# Planemo: a command-line toolkit for developing, deploying, and executing scientific data analyses

**DOI:** 10.1101/2022.03.13.483965

**Authors:** Simon Bray, Matthias Bernt, Nicola Soranzo, Marius van den Beek, Bérénice Batut, Helena Rasche, Martin Čech, Peter Cock, Anton Nekrutenko, Björn Grüning, John Chilton

## Abstract

There are thousands of well-maintained high-quality open-source software utilities for all aspects of scientific data analysis. For over a decade, the Galaxy Project has been providing computational infrastructure and a unified user interface for these tools to make them accessible to a wide range of researchers. In order to streamline the process of integrating tools and constructing workflows as much as possible, we have developed Planemo, a software development kit for tool and workflow developers and Galaxy power users. Here we outline Planemo’s implementation and describe its broad range of functionality for designing, testing and executing Galaxy tools, workflows and training material. In addition, we discuss the philosophy underlying Galaxy tool and workflow development, and how Planemo encourages the use of development best practices, such as test-driven development, by its users, including those who are not professional software developers. Planemo is a mature project widely used within the Galaxy community which has been downloaded over 80,000 times.

## Introduction

The Galaxy project provides web browser access to command-line scientific software, together with the necessary compute resources, in a convenient, shareable and reproducible way, to researchers around the world [1]. Over eight thousand tools are available for installation onto any Galaxy server; users can run these individually, connect multiple tools together to form workflows, and finally perform complex analyses, without the need to access a command line. While Galaxy itself does not require any significant computational skills to use, development and maintenance of new tools and workflows benefit from sophisticated infrastructure with both human and automated components. The process of integrating software into Galaxy requires knowledge of both the command-line interface of the underlying software and the schema used by Galaxy to define tools, in order to be able to write a ‘Galaxy tool wrapper’ mapping dataset inputs, parameter inputs and outputs between them. Once written, wrappers, as well as other Galaxy artifacts such as workflows or training material [2], are amenable to routine processes such as testing, deployment and regular updates, all of which can be automated using continuous integration (CI) systems. Here we present Planemo, a versatile library and command line application which is used extensively as a software development kit by Galaxy or Common Workflow Language (CWL) [3] tool, workflow and training material developers, and as a toolkit for Galaxy ‘power users’. Planemo provides a simple but powerful command-line interface for tool and workflow development and deployment, which encourages and enforces good practices for software development. In addition, it enables automated deployment of developed tools and automatic updates of the software dependencies used internally by each Galaxy tool. The testing functionality included in Planemo has been successfully integrated into CI workflows of the major tool and workflow repositories, which helps to ensure the creation of high quality tool wrappers and workflows.

Planemo is structured into numerous subcommands, which provide a broad range of functionality. Here we discuss a selection of the most important functionalities, grouped around the following themes: 1) development of Galaxy tools, workflows, tutorials, and CWL tools; 2) deployment of the developed tools and workflows; 3) automated tool and workflow dependency updates and 4) tool and workflow execution. Table 1 summarizes this functionality, and Fig. 1 provides a graphical overview. In addition to its use as a command-line application, Planemo can also be used as a library by other projects. An example is the Planemo Training Development Kit project (https://github.com/galaxyproject/ptdk), which provides Planemo’s functionality for creating training material for Galaxy workflows via a webserver.

**Table 1.**
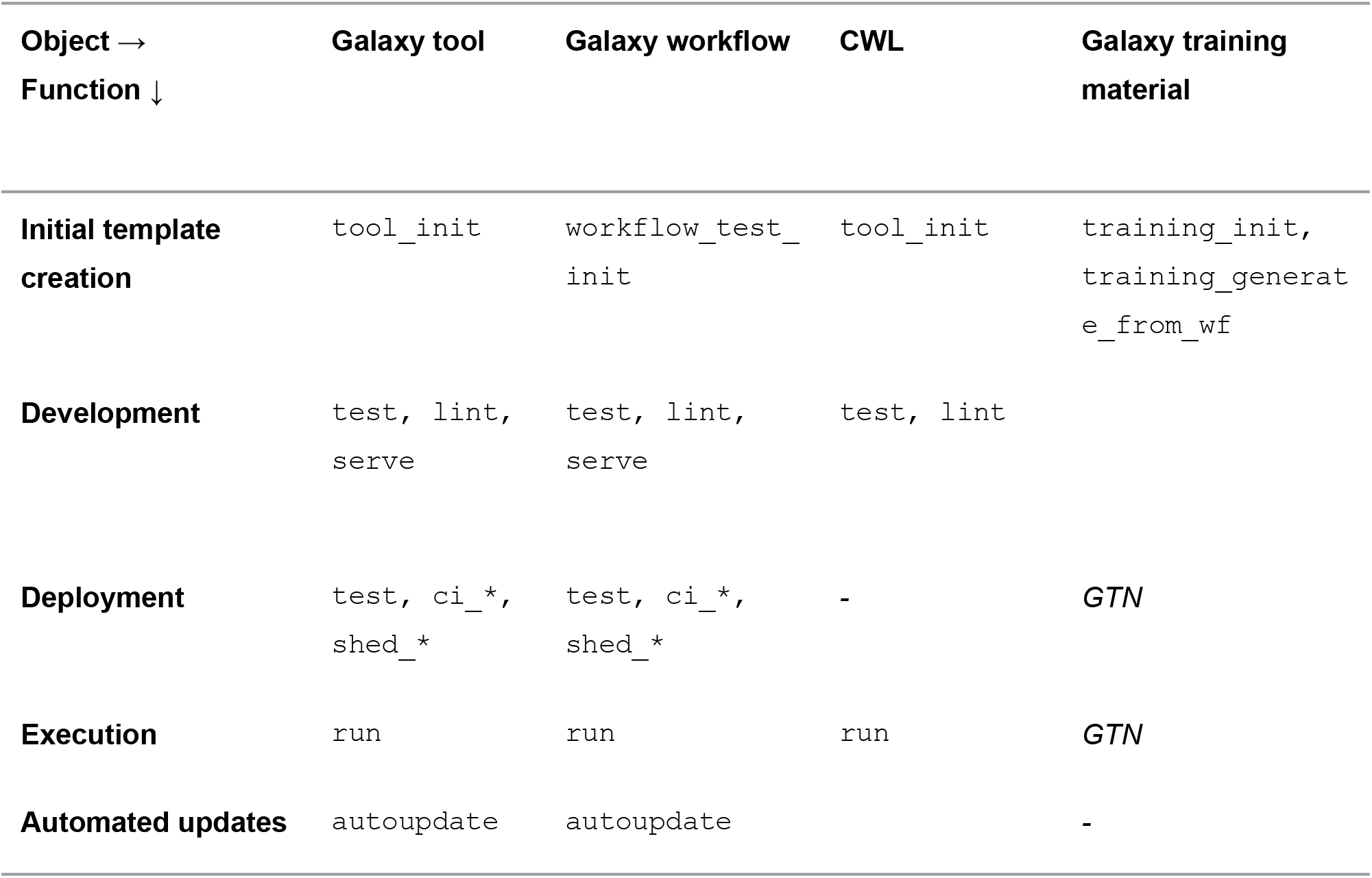
Overview of Planemo functionality and subcommands. Columns represent artifacts that can be created or manipulated with Planemo, rows represent different actions that can be performed on them. Italics represent actions which are performed without using Planemo: trainings are deployed using Jekyll and executed by users following the training material in the graphical interface.

| Object →<br>Function ↓ | Galaxy tool | Galaxy workflow | CWL | Galaxy training material |
| --- | --- | --- | --- | --- |
| **Initial template creation** | tool_init | workflow_test_init | tool_init | training_init, training_generate_from_wf |
| **Development** | test, lint, serve | test, lint, serve | test, lint |  |
| **Deployment** | test, ci_*, shed_* | test, ci_*, shed_* | - | *GTN* |
| **Execution** | run | run | run | *GTN* |
| **Automated updates** | autoupdate | autoupdate |  | - |

**Figure 1.**
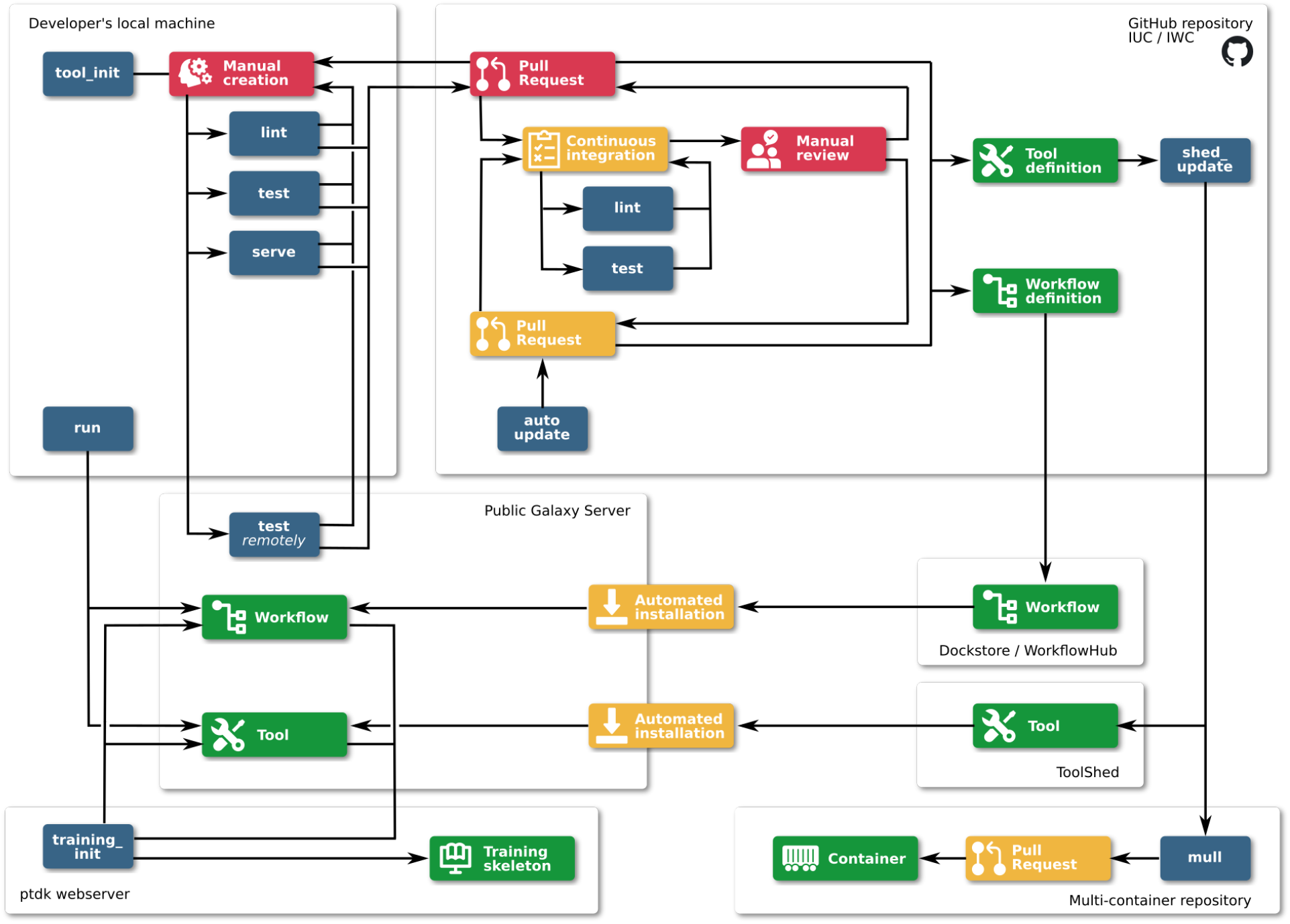
Overview of the use of Planemo for development, deployment, and execution of Galaxy tools, workflows and training materials. Red = manual work, blue = Planemo commands, yellow = automated steps, green = created artifacts.

## Methods

### Software design

Planemo is implemented as a Python package and distributed via GitHub, PyPI and Bioconda [4]. As already described in the Introduction, Planemo is a highly flexible, multifunctional software, which can be used for: 1) different types of artifacts (e.g. tools, workflows), 2) different workflow/tool languages and management systems (e.g. Galaxy, CWL), 3) different tasks (e.g. linting, testing, executing). To handle this variety, Planemo defines two central abstractions: Runnables and Engines. Runnables include tools and workflows written for either Galaxy or CWL; an Engine provides access to an external piece of software (such as Toil or Galaxy) capable of executing a particular Runnable. Each Engine has various methods (e.g. run(), test()), which define a particular interaction with a Runnable.

Engines are provided for both local and external Galaxy servers, as well as for cwltool [5] and Toil [6]. These interact with their respective workflow management systems via the cwltool and Toil Python modules (for CWL), and via the BioBlend library [7], which provides access to the Galaxy API through Python. Numerous lower-level functions and classes are provided to connect the Engines with the underlying functionality.

Some tasks cannot be easily described in the context of these abstractions; for example, linting of tool or workflow definitions requires only that the structured document containing the definition be compared with a schema. Other examples include the functionality for automatic updates of software dependencies and generation of training material. Planemo handles these cases using separate classes and functions.

Planemo is most frequently used as a command-line application, using a command-line interface written using the Click package to provide a straightforward way to access the components described above. Multiple subcommands expose some of the most important tasks a user might want to perform. For example, a user could run `‘planemo test tool.xml’` to test a Galaxy tool wrapper. Planemo will detect the type of Runnable (Galaxy tool) represented by the filepath and start the appropriate Engine (temporary local Galaxy instance), execute the Runnable on it, collect the results, and compare them to predefined test data to determine a pass or fail status. All subcommands can be configured by appending flags and options.

### Implementation of continuous integration jobs

While Planemo is designed primarily with developers and users in mind, commands often need to be executed as part of automated continuous integration (CI) jobs – for example, testing of newly created Galaxy tools after submission to a GitHub repository. Galaxy tools and workflows are hosted over multiple repositories; to ensure a unified approach to testing, a GitHub CI action is provided. The CI workflow consists of the following components:

1. Identifying modified tools and repositories using `‘planemo ci_find_repos’` and `‘planemo ci_find_tools‘`.

2. Linting of Galaxy tools using `‘planemo lint’`.

3. Testing the tools – as this is the most time-consuming step, the tools found are chunked and multiple jobs run in parallel.

4. Linting of Python and R scripts packaged together with the tools.

5. If the PR is approved and merged: deployment to the Toolshed with ‘planemo shed_update’.

### Definition of terms

Planemo’s features rely on and are interdependent with a variety of other subprojects within and related to the Galaxy community. We therefore first outline a few of these.

#### IUC

The Intergalactic Utilities Commission [8] maintains a central repository of Galaxy tool wrappers, currently hosted on GitHub. New wrappers are added by means of a GitHub pull request, reviewed by IUC members, and are tested by automated CI. After approval, the tool is automatically deployed to the Galaxy ToolShed. Tools are subject to further automatic updates, as new versions of software dependencies are released. The IUC serves as a model for smaller communities developing wrappers for more specialized tools (for example, Galaxy-P [9] for proteomics) and has developed a set of guidelines for tool development.

#### Bioconda/BioContainers

Each Galaxy tool has certain dependencies, which are typically installed either using the Conda package manager [10] or within a container (Docker [11] or Singularity [12]). Development and maintenance of the necessary Conda packages or containers is performed by the Bioconda and Biocontainers [14] communities, which collaborate closely with the Galaxy project.

#### ToolShed

A central ‘app store’ for Galaxy tools Any user can upload to the ToolShed [13], but most high-quality tools are developed collaboratively on an open platform like GitHub (for example by the IUC) and deployed automatically.

#### IWC

maintains a set of curated workflows [14], consisting of multiple component Galaxy tools, which are hosted on GitHub and deployed to Dockstore [15] and the Workflow Hub [16], analogously to the development and deployment of Galaxy tools to the ToolShed by the IUC.

#### Galaxy Training Network

A repository for tutorials, each describing a method for data analysis in Galaxy [1]. Each tutorial is made up of multiple steps and therefore corresponds to a Galaxy workflow, which forms the skeleton around which the tutorial is built.

#### Continuous Integration (Workflow)

A workflow run remotely on a build server which tests and deploys Galaxy artifacts developed. It should not be confused with a Galaxy workflow.

#### Tool

Artifact defined by a tool wrapper and stored in the ToolShed, allowing users to access the functionality of the underlying software via Galaxy.

#### Galaxy Tool Wrapper

Structured document defining a Galaxy tool; it maps dataset inputs and outputs and other parameters between the underlying command-line tool and the Galaxy API.

#### Galaxy Workflow

a directed acyclic graph in which nodes can be dataset inputs or outputs, parameter inputs, or tools. More informally, a combination of multiple individual tools into a single pipeline, which once assembled can be executed as if it were a single tool.

#### Collection

a group of individual datasets linked together in a directory-like structure. When a tool is run on a collection, individual jobs are generated for each of the datasets which make up the collection. In combination with workflows, collections allow Galaxy users to scale up analyses to deal with large sets of data.

### Documentation

Planemo’s documentation is hosted on a ReadTheDocs site: https://planemo.readthedocs.io. In addition, several tutorials are available as part of the Galaxy Training Network:

- Creating Galaxy tools from Conda through deployment: https://training.galaxyproject.org/training-material/topics/dev/tutorials/tool-from-scratch/tutorial.html
- Creating training material with Planemo: https://training.galaxyproject.org/training-material/topics/contributing/tutorials/create-new-tutorial/tutorial.html
- Automating Galaxy workflows using the command line: https://training.galaxyproject.org/training-material/topics/galaxy-interface/tutorials/workflow-automation/tutorial.html
- Test-driven development with Planemo: https://planemo.readthedocs.io/en/latest/writing_advanced.html#test-driven-development

## Results and Discussion

### Galaxy tool development

A Galaxy tool is defined by a wrapper for an underlying software (or code), which maps its dataset inputs, parameter inputs and outputs to a command-line script executed by Galaxy. When running a tool in the Galaxy interface, a user selects their preferred choices for the exposed dataset and parameter inputs. The Galaxy server then constructs the command, schedules it as a job onto appropriate compute resources, collects the results once the job has completed, and returns them to the user.

Writing Galaxy tool wrappers requires a thorough knowledge of the underlying software and also an understanding of the Galaxy tool schema which defines how Galaxy wrappers are written. The tool schema is defined in a simple manner, in order to make the process of wrapping software as accessible as possible [17]. Planemo provides several helpful features which assist tool developers in creating high-quality wrappers that meet community-defined standards, such as those [18] developed by the Intergalactic Utility Commission (IUC). These features are implemented as subcommands, e.g. ‘`planemo test`’. Planemo also helps to enforce software development best practices such as writing tests for all tools and linting the wrapper definitions to avoid bugs and ensure a coherent and readable style. Further support for tool development standards is provided by the Galaxy Language Server [19], an implementation of the Language Server Protocol [20] and a Visual Studio Code extension for Galaxy tools, which can be used side-by-side with Planemo.

A common starting point for tool development is the ‘`tool_init`’ subcommand. To use this, the developer provides a variety of options, including an example command line, tool name, inputs, outputs and software requirements, from which Planemo generates a skeleton tool wrapper. Most of the ‘`tool_init`‘ parameters are optional, but the more that are provided, the more detailed the initial skeleton will be.

The developer can then inspect and edit the generated file, adding more parameters and increasing the complexity of the wrapper logic by incorporating conditionals and repeat elements if necessary. As they continue to edit, they can use the ‘`lint`‘ subcommand to validate the wrapper under development. Planemo’s linting forces wrappers to match Galaxy’s tool schema, ensuring stylistic consistency and preventing some errors such as mismatched file formats. Crucially, Planemo recommends that wrappers define at least one test case to ensure the development of high-quality, portable, reliable and functional tools, and this recommendation is strictly enforced by the IUC’s and other tool repositories. Once tests are defined, together with an initial tool definition, the developer can start to run the tests using the ‘`test`‘ subcommand. This launches a transient Galaxy server on the developer’s computer, installs the Galaxy tool under development, together with all software dependencies, and executes the tests specified within the tool wrapper. The results of the tests are then returned to the developer, by default using a report defined using JSON and HTML, although other format types are also supported (xUnit, jUnit, Markdown and Allure).

Planemo encourages the use of test-driven development [21], a software development principle which states test cases should be written before a new feature is developed. Test-driven development is an industry-wide best practice. Defining extensive test cases at the start of the process covering the required features provides a focus for development, and results in more robust and better documented code containing fewer bugs. The tool developer is forced to adopt the perspective of the Galaxy user from the start to consider possible use-cases of the software for which tests need to be written. Initial test failures lead to iterative refinement of the wrapper, until a fully-functional Galaxy tool, which passes all tests, is produced.

Once tests are passing, the developer should optimize the tool interface which is presented to the user of the tool. To facilitate this, Planemo provides the ‘`serve`‘ subcommand, which launches a Galaxy server with the new tool installed, allowing the developer to inspect the rendering of the wrapper in the graphical interface and to perform manual testing. The developer should also improve the documentation of the tool, by annotating each of the tool parameters, as well as writing a help section to explain the tool’s aim and usage, which appears beneath the tool parameters in the graphical interface.

### Common Workflow Language tool development

In addition to Galaxy tools, Planemo also acts as a software development kit for CWL tools. The same subcommands described can be used for this purpose, including ‘`tool_init`’ and ‘`test`’. By appending the ‘`--cwl`’ argument to the ‘`tool_init`’ subcommand, Planemo generates a template for a CWL tool definition, rather than a Galaxy wrapper. The test and lint commands then detect that the input file is a CWL wrapper and process it accordingly. Tools are tested by executing with the CWL engine cwltool and comparing the result with test data or specified assertions, in the same way as for Galaxy tools. The completed wrapper can be run using any CWL engine, such as cwltool, Toil, Arvados [22] or Galaxy.

### Galaxy workflow development

Workflows are created in Galaxy by connecting together multiple tools (i.e. an output of one tool becomes an input for the following one) in order to automate complex analyses. Unlike tools, workflows can be defined and edited in Galaxy’s graphical workflow editor; often the starting point is an interactive analysis (a Galaxy history) from which a workflow can be extracted automatically. It is also possible to manually author workflows in the gxformat2 workflow language [23], and the user can switch between manually writing workflows and editing in the graphical interface using the ‘`workflow_edit`’ subcommand, which spins up a Galaxy instance with the workflow under development pre-installed for editing. Planemo additionally facilitates the creation of test cases by providing the option of generating them automatically from a pre-existing workflow invocation.

Once a draft version of the workflow exists, it should be iteratively improved in the same way as for tools, using the same lint, test and serve subcommands already introduced. The ‘`workflow_lint`‘ subcommand checks workflows for errors and conformance with best practices—a command-line interface mirroring functionality which is also provided by the Galaxy graphical workflow editor. For example, workflows which are missing test cases, labeled outputs, or essential metadata fail linting. Running the ‘`test`‘ subcommand launches a local Galaxy instance, installs the tools used in the workflow, uploads the workflow and executes it on the provided input test data. In the same way as for tool testing, the workflow outputs are downloaded and compared to the test data, resulting in either a pass or fail status. In some cases, it can be convenient to run testing on an existing public server, such as https://usegalaxy.org, https://usegalaxy.eu, or https://usegalaxy.org.au; this is also supported by Planemo. Running the ‘`serve`‘ subcommand provides a local Galaxy server with the workflow and the needed tools pre-installed, which can be used for workflow development and fine-tuning.

### The philosophy of Galaxy tool and workflow development

After the previous discussion of the process of tool and workflow development, the question arises how software complexity should be divided between the tool and the workflow level. Should most of the effort go into developing workflows, keeping tools as simple as possible and flexibly rewrapping the underlying software depending on the demands of a particular workflow, or should developers invest time creating complex and multifunctional tools which can be reused without modification in multiple workflows?

Galaxy leans heavily towards the second of these two options, as does CWL, though the following discussion will focus on Galaxy. Galaxy encourages the creation of modular tools which are usable in isolation, so they can be used interchangeably in multiple different workflows. Tools generally encapsulate most of the complexity of the underlying software, allowing workflows to be simply constructed in a graphical interface by connecting the component tools. Workflows can thus be thought of as complex structures built from the same fundamental building blocks, which can be constructed without knowledge of the internal functionality of the individual tools. This has several advantages with regard to the user experience: building workflows becomes a far less daunting task, and tools can also be used individually in the graphical interface, which makes Galaxy accessible to new users and enables its use as a teaching environment for scientific analysis.

Another advantage of this approach is the “separation of concerns”, a design principle in computer science. Different groups of scientists can develop and apply specialized and complementary areas of knowledge: the tool developer can concentrate on describing and developing the Galaxy tool, without considering any downstream workflows that will be created later. On the other hand, the workflow developer can construct complex, high-level pipelines, without the detailed understanding of the component tools and the command-line possessed by the tool developer. This has the dual advantage that workflows can be treated on a more abstract level and that the workflow creation process is made accessible for a far greater number of users.

Separation of concerns between tools and workflows also benefits security. Executing untrusted software on a compute cluster is highly undesirable; thus workflows need to be assessed for security risks before execution. For many workflow management systems, this assessment must be repeated for each workflow. By contrast, as the Galaxy tool review process involves checking tools for security issues before merging, a system administrator can deploy tools developed by the IUC or similar high-trust communities with confidence. The question of workflow security is thus made redundant: if the component tools are trusted, a workflow based on those tools can likewise be trusted.

These advantages must be balanced against the time investment required from community members to build up a diverse set of tools, to allow the construction of scientifically interesting workflows. Nonetheless, the Galaxy community, facilitated by Planemo, has succeeded in developing such a toolset and making it available to the scientific community.

### Continuous integration for community repositories

Galaxy has a large and vibrant community of tool and workflow developers, creating Galaxy tools in a wide range of scientific fields, ranging from genomics to proteomics, computational chemistry and climate science. As a result, a large number of high quality tools already exist and are actively maintained over several GitHub repositories, centered around the main IUC repository; the IWC (see Methods for definition) performs the equivalent function of a repository for Galaxy workflows. Building these communities has required many years of work by multiple contributors; in order to streamline the process and ease the burden on the tool developers, developing infrastructure to facilitate human review and automate as much as possible is essential. Planemo forms the core of this infrastructure.

Once a developer has completed the tool wrapper or workflow, they can submit it to a community repository, usually hosted on GitHub, for review. Alternatively, they may also deploy it themselves (for example, to the ToolShed or WorkflowHub), but submission to a community repository is encouraged to ensure the code is thoroughly reviewed and to publicize the new tool or workflow. Community repositories are configured to run the linting and testing checks already described after submission, via a continuous integration (CI) workflow. Planemo provides a couple of simple subcommands, ‘`ci_find_repos`‘ and ‘`ci_find_tools`‘, to identify tools which have been added or modified. Both of these allow chunking of tools in order to parallelize the testing process over multiple CI jobs. As part of the CI testing, linting and testing of the tools is repeated, as well as linting of any Python and R scripts added together with the new tool wrappers. These steps ensure the submitted tools are of high quality, enforce consistent standards on the code and reduce the maintenance burden for the entire community.

If all tests pass and the proposed new tool or workflow is accepted by the community, another CI job is initiated to deploy it to the ToolShed. This makes use of Planemo’s ‘`shed_update`‘ command, which uses the ToolShed credentials associated with the repository to upload the newly created tool. Once it is available on the ToolShed, it can easily be installed onto any Galaxy server.

The entire process, consisting of automated testing, human review and automated deployment, ensures the creation of high-quality, trustworthy tools which can be safely installed and used. It requires several more specialized steps, which go beyond the simple Planemo subcommands that the developer runs on their local machine. To package these CI workflows into a single unit, a GitHub Action is provided [24] which can be reused in other tool repositories. New tool repositories with the same structure as the IUC repository can be conveniently created from a template repository created by the Galaxy community [25].

### Automation of tool and workflow updates

Another feature offered by Planemo is automatic updates of Galaxy tool and workflow software dependencies, using the `‘autoupdate‘` subcommand. In combination with separate autoupdate features already developed by the Bioconda and conda-forge [26] communities, this forms a sequence of semi-automated software update procedures, which are triggered by an official release of new source code. After this new release appears, this chain ensures that new Conda packages, new Docker and Singularity containers, updated Galaxy tools and finally updated Galaxy workflows are generated (Fig. 2). At each step, a CI job detects the artifact published in the previous step and initiates the process of updating a dependent artifact, generally by means of a GitHub pull request (PR).

**Figure 2.**
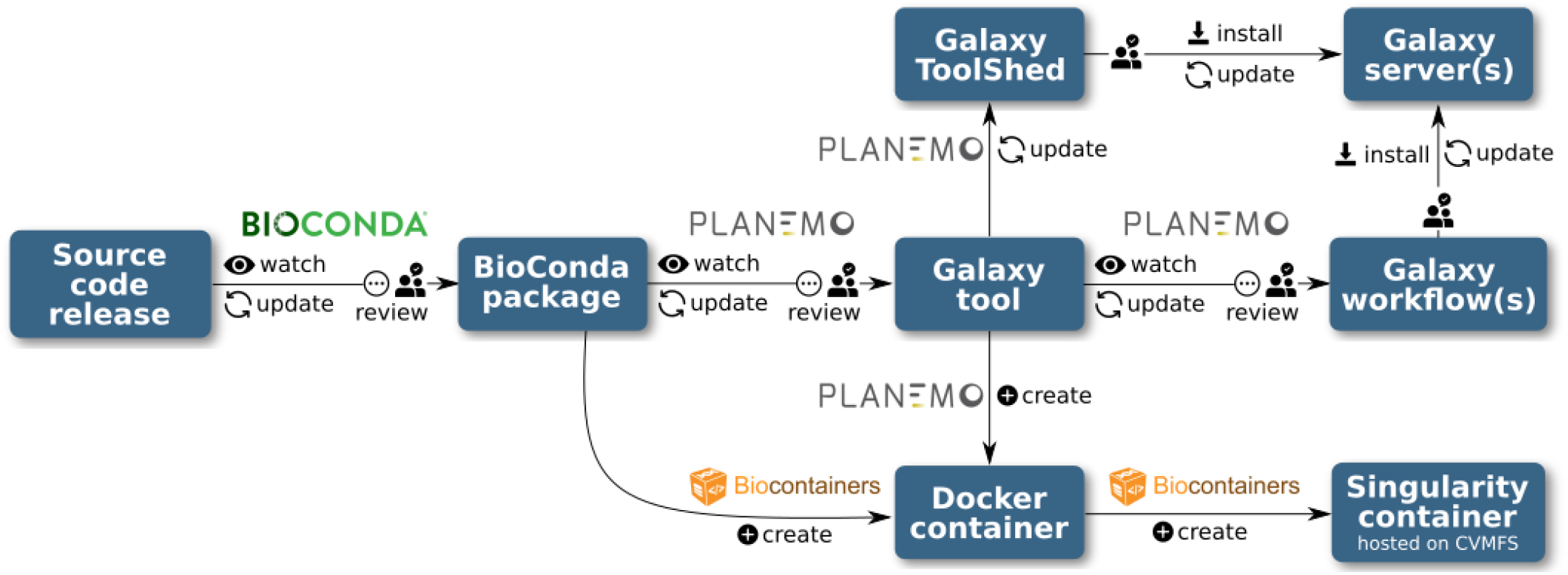
Automation pipeline for Bioconda packages, BioContainers, Galaxy tools and workflows. Steps marked in red require human review; steps marked in blue are fully automated.

The CI pipelines developed by Bioconda and conda-forge monitor the Conda recipes they maintain, regularly checking the links provided in the recipes for new releases. When the developers of an upstream software package release a new version, the CI creates a PR to update the package recipe. Once the PR is reviewed and merged, newly built packages are uploaded to the Anaconda repository.

In parallel, a bot [29] running the `‘autoupdate‘` subcommand monitors the Galaxy tool wrappers maintained by the IUC, as well as a few other smaller communities, checking the dependencies defined in the tool wrapper. Once an updated Bioconda or conda-forge package is published in the step above, the Planemo autoupdate bot detects this and updates the dependencies section of the Galaxy tool accordingly. A PR is then submitted to the GitHub repository, to be reviewed and manually updated if necessary, before it is merged and deployed as described in the “CI for community repositories” section.

Galaxy tools can specify multiple dependencies. If these dependencies are installed via Conda, the packages can be simply installed into a single environment, but if dependency installation is achieved using containers, a new container must be built for each required combination of dependencies. This is achieved by the ‘mulled build’ infrastructure; a CI job triggers the building of a Docker container for each new combination of packages, on publication of new Galaxy tool versions. Another CI job is responsible for generating Singularity containers from the new Docker containers, which are made available by the BioContainers and Galaxy communities via a CernVM file system (CVMFS) [27]. These steps do not require manual review.

The Planemo autoupdate bot also monitors the Galaxy workflows maintained by the IWC and checks whether new versions exist for each of the component tools. Once a new tool version is created (either by the upstream tool autoupdate step, or a tool developer), the workflow definition file hosted by the IWC is modified accordingly and a PR submitted for review (Fig. 3).

**Figure 3.**
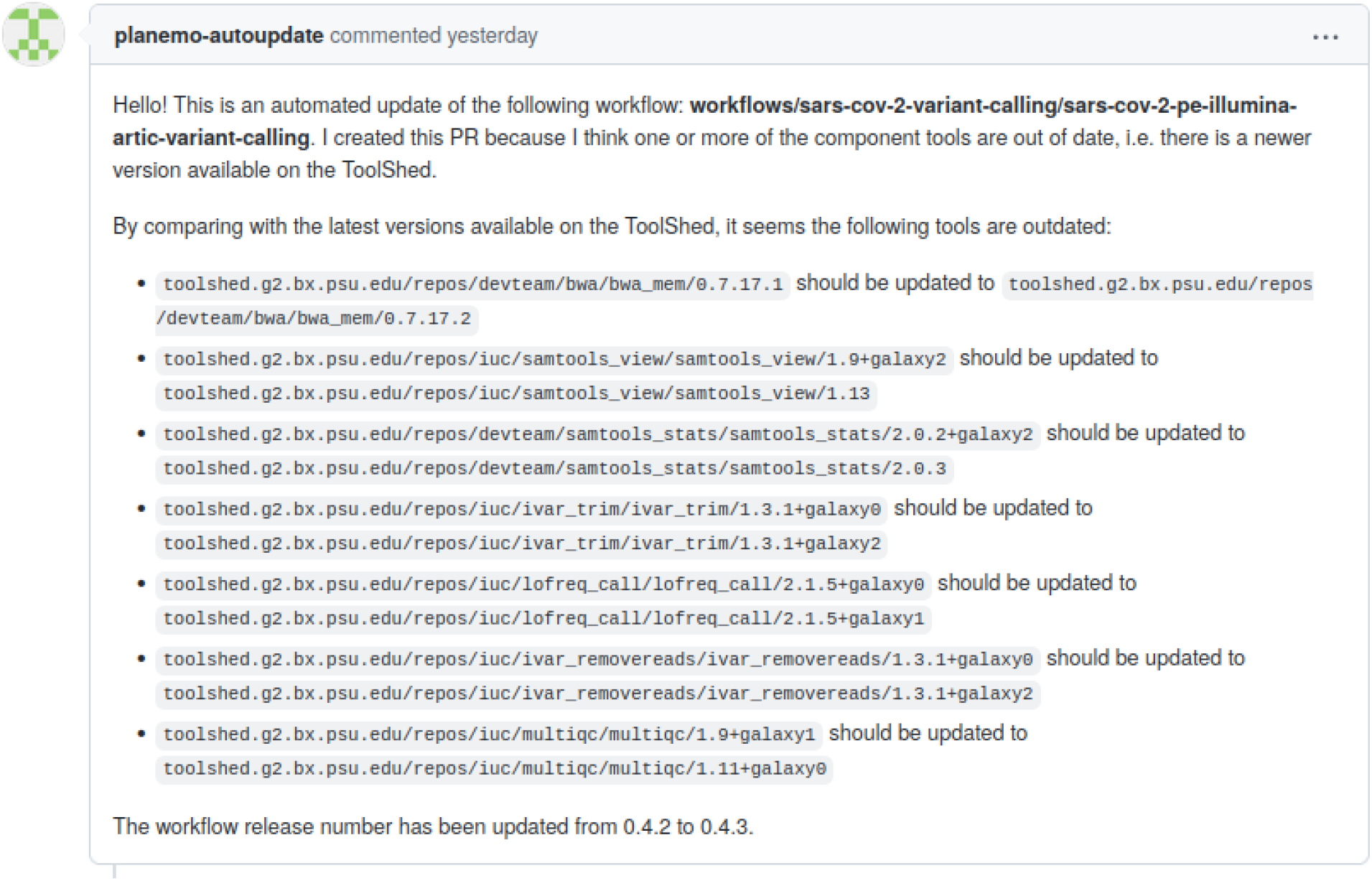
An example GitHub pull request created by the Planemo autoupdate bot, updating a workflow hosted on the IWC.

### Execution

Apart from providing assistance with tool and workflow development and deployment, Planemo is also a useful resource for Galaxy power users who need to launch high-throughput data analyses. Galaxy is traditionally accessed via a graphical interface in the web browser, and features such as Galaxy collections already provide a high level of parallelization to users of the graphical interface. Nonetheless, there are important scenarios in which a user might need to run individual workflows hundreds or thousands of times, in which the data cannot be grouped into collections ahead of time—for example, for variant calling of SARS-CoV-2 genomic data, in which a huge amount of new data is published continuously [28]. As a convenient alternative to the graphical interface, Planemo allows workflow execution to be scheduled programmatically using the ‘`run`‘ subcommand, either on a local machine or a larger Galaxy server. ‘`planemo run`‘ can be embedded in scripts of varying complexity, which can be scheduled and controlled via CI systems or message queues to run workflows on demand - such as on new data appearing or tool updates.

Internally, Planemo executes workflows by submitting them to the chosen server via Galaxy’s API. Requests to the API are made using BioBlend, a library which wraps many API endpoints as Python methods. It is also possible to execute workflows directly using BioBlend, or simply by making API calls using a tool such as cURL. While this approach does offer a high level of flexibility, it requires the user to possess a high level of knowledge of the API (for example, the correct format to submit workflow parameters) and often requires the creation of custom scripts. By contrast, Planemo’s ‘run‘ subcommand offers a high-level interface to execute workflows, monitor them during execution, and report on their status after completion, packaged as a single command.

For tool and workflow development, the artifacts under development are generally tested against an ephemeral local Galaxy instance, which is deleted after use. While this is also supported by the ‘`run`‘ subcommand, with the workflow outputs saved to a specified location, this approach is not scalable for workflows which demand long compute times, with large data inputs, or with workflows which need to be executed multiple times. In many cases, the user may prefer to make use of established, stable infrastructures, such as a public Galaxy instance or a private instance administered by their research group. Planemo allows external Galaxy instances to be specified for all ‘`run`‘ and ‘`test`‘ commands by providing the server URL and user API authentication key on the command line. As it is inconvenient and insecure to enter the API key with each command, Planemo also allows users to define profiles, in which the URL and API key is configured for each server. The user can then define multiple profiles and run workflows on different servers simply by appending, e.g. ‘`--profile usegalaxy-org`‘ or ‘`--profile private-server`‘ to the command.

Planemo provides numerous command line options to configure the workflow execution process. The name of the history in which the new invocation is created, as well as a list of Galaxy tags to add, can be specified via the command line. In addition, Planemo and Galaxy allow both datasets and workflows to be specified via hexadecimal IDs which point towards a Galaxy object on an external server, rather than by referring to a local path. This has the advantage of avoiding multiple uploads of the same dataset or workflow, if the workflow has to be executed multiple times. Planemo can also be configured to either wait until the workflow has completed, and download the output datasets created, or to terminate once the workflow has been successfully scheduled. In the latter case, the ‘`list_invocations`‘ command can be used to monitor running workflows and to return the number of jobs which have succeeded, failed, or incomplete. If jobs have failed—for example, due to transient server issues— the user can also choose to restart them using the ‘rerun‘ subcommand.

### Training material

Planemo provides utilities for developing tutorials for different types of data analysis with Galaxy. The Galaxy Training Network, accessible via https://training.galaxyproject.org, provides a range of training material including slide decks, tutorials and videos. In particular, the tutorials are written in Markdown and rendered using Jekyll, and often feature ‘hands-on boxes’ which describe the exact combination of parameters and input which users need to submit when running a Galaxy tool. Most tutorials instruct the trainees to run several Galaxy tools in sequence, and thus correspond to a Galaxy workflow.

Planemo provides two subcommands, `‘training_init‘` and `‘training_generate_from_wf‘`, which generate a directory structure for a new tutorial, containing skeleton Markdown files defining the tutorials. These files already contain sections and hands-on boxes for each tool, with the tool inputs and parameters predefined, ensuring a high level of consistency in the appearance and quality of the tutorials produced. The training developer can then take these templates and expand them with additional information, questions, diagrams and citations to produce the completed training. They also need to provide input datasets, which are usually stored on Zenodo. To populate a Galaxy server with these datasets, the training developer should also provide a data library file, which can be generated using the `‘training_fill_data_library‘` subcommand, including the Zenodo links and file formats of the datasets.

A major aim of the Galaxy Training Network project is improving accessibility for new contributors, including for scientists who are not comfortable with command-line software. As a result, the Planemo functionality relating to training material development is provided in webserver form as the Planemo Training Development Kit (PTDK). The application is written using Flask and deployed with Heroku; it can be accessed via https://ptdk.herokuapp.com. The interface allows the selection of the same options as the Planemo commands, with the additional option of specifying a workflow for generating the training using its ID from one of the major public Galaxy servers.

## Conclusion

We have presented Planemo, a library and application which has already achieved widespread usage among Galaxy tool, workflow and training material developers, Galaxy power users, and as part of numerous automated deployment solutions. Planemo provides the developers of command-line software with an easy way to create a graphical interface, taking advantage of the many features developed by the Galaxy community and the compute resources provided by public Galaxy instances. We have described the complex infrastructure the Galaxy community has developed for creating and interacting with artifacts such as tools, workflows and training material. Planemo plays the crucial role of bridging the gaps between the human and automated components of this infrastructure, freeing members of the community to devote their time to developing, reviewing and performing novel scientific analyses.

## Acknowledgements

The authors are grateful to the broader Galaxy community for their support and software development efforts. This work is funded by NIH Grants U41 HG006620 and NSF ABI Grant 1661497. Usegalaxy.eu is supported by the German Federal Ministry of Education and Research grants 031L0101C and de.NBI-epi to BG. Usegalaxy.org.au is supported by Bioplatforms Australia and the Australian Research Data Commons through funding from the Australian Government National Collaborative Research Infrastructure Strategy. The funders had no role in study design, data collection and analysis, decision to publish, or preparation of the manuscript.

